# A brain-inspired algorithm improves “cocktail party” listening for individuals with hearing loss

**DOI:** 10.1101/2024.05.01.592078

**Authors:** Alex Boyd, Virginia Best, Kamal Sen

## Abstract

Selective listening in competing-talker situations (restaurants, parties, etc.) is an extraordinarily difficult task for many people. For individuals with hearing loss, this difficulty can be so extreme that it seriously impedes communication and participation in daily life. Directional filtering is one of the only proven ways to improve speech understanding in competition, and most hearing devices now incorporate some kind of directional technology, although real-world benefits are modest, and many approaches fail in competing-talker situations. We recently developed a biologically inspired algorithm that is capable of very narrow spatial tuning and can isolate one talker from a mixture of talkers. The algorithm is based on a hierarchical network model of the auditory system, in which binaural sound inputs drive populations of neurons tuned to specific spatial locations and frequencies, and the spiking responses of neurons in the output layer are reconstructed into audible waveforms. Here we evaluated the algorithm in a group of adults with sensorineural hearing loss, using a challenging competing-talker task. The biologically inspired algorithm led to robust intelligibility gains under conditions in which a standard beamforming approach failed. The results provide compelling support for the potential benefits of biologically inspired algorithms for assisting individuals with hearing loss in “cocktail party” situations.

## INTRODUCTION

One of the most challenging listening tasks encountered by people in their daily life is to understand what a talker of interest is saying in an acoustically cluttered environment, especially one that contains other people talking at the same time^1,2^ (the proverbial “cocktail party problem”). Listeners with sensorineural hearing loss (including older people with typical age-related declines in hearing) experience extreme difficulties in these situations, and in many cases ease-of-communication and willingness to attend social gatherings is seriously impeded.^3,4^ We are now beginning to understand the devastating longer-term consequences of untreated hearing difficulties, which include declines in cognitive health, and the associated societal and economic costs.^5^

One proven way to improve speech understanding in noise is via directional filtering.^6^ Directional microphones are by now almost ubiquitous in hearing aids and provide broad filtering that attenuates sounds arising from behind the listener to improve the signal-to-noise ratio (SNR) for those in front. Narrower tuning can be achieved by using larger numbers of microphones^7^ (e.g., arranged in an array on a headband or eyeglasses). A number of two-channel (binaural) beamforming algorithms achieve narrow tuning by combining the signals at the left and right ears.^8^ One issue with most previous solutions is that they sacrifice natural spatial information, which can counteract any improvements in SNR they provide, especially in complex situations with competing talkers.^9,10^ Deep neural network approaches to sound segregation have made impressive leaps in recent years and under the right conditions can effectively isolate a sound of interest from a complex mixture.^11,12^ These approaches, however, are computationally expensive and not yet well-suited for low-power real-time applications.^13^

We recently developed a biologically oriented sound segregation algorithm^14^ (BOSSA), which is designed to separate competing sounds based on differences in spatial location. Taking its inspiration from binaural auditory system, this algorithm requires only two input signals, and does not sacrifice spatial cues. It is also well-suited for low-power real-time applications and thus could have real utility in wearable hearing assistive devices. BOSSA was developed and optimized using objective intelligibility measures, and has been evaluated behaviorally in a group of young listeners with audiometrically normal hearing.^14^ In this population, BOSSA provided robust improvements in the intelligibility of a target talker embedded in a challenging speech mixture. To date, BOSSA has not been evaluated in the population who most stand to benefit from assistance in cocktail party situations, which is individuals with hearing loss.

In the current study, we recruited adults with bilateral sensorineural hearing loss, and measured the benefits provided by BOSSA for the task of understanding one talker in a mixture of five competing talkers. We compared the performance of BOSSA to a binaural implementation of the current industry standard beamforming approach (minimum variance distortionless response, or MVDR) employed in hearing aids.^8^

## RESULTS

Figure 1 shows the average proportion of words correctly identified as a function of target-masker ratio (TMR) for four different multitalker scenarios (panels A-D), and for four different sound processing conditions (colored lines). These functions show that performance improved systematically with TMR, as expected, but also differed systematically between processing conditions. In each scenario,, performance for two different versions of BOSSA (DiffMask and RatioMask) was better than performance for the Natural (unprocessed) condition, and was also better than for the standard MVDR beamformer condition.

**Figure 1.**
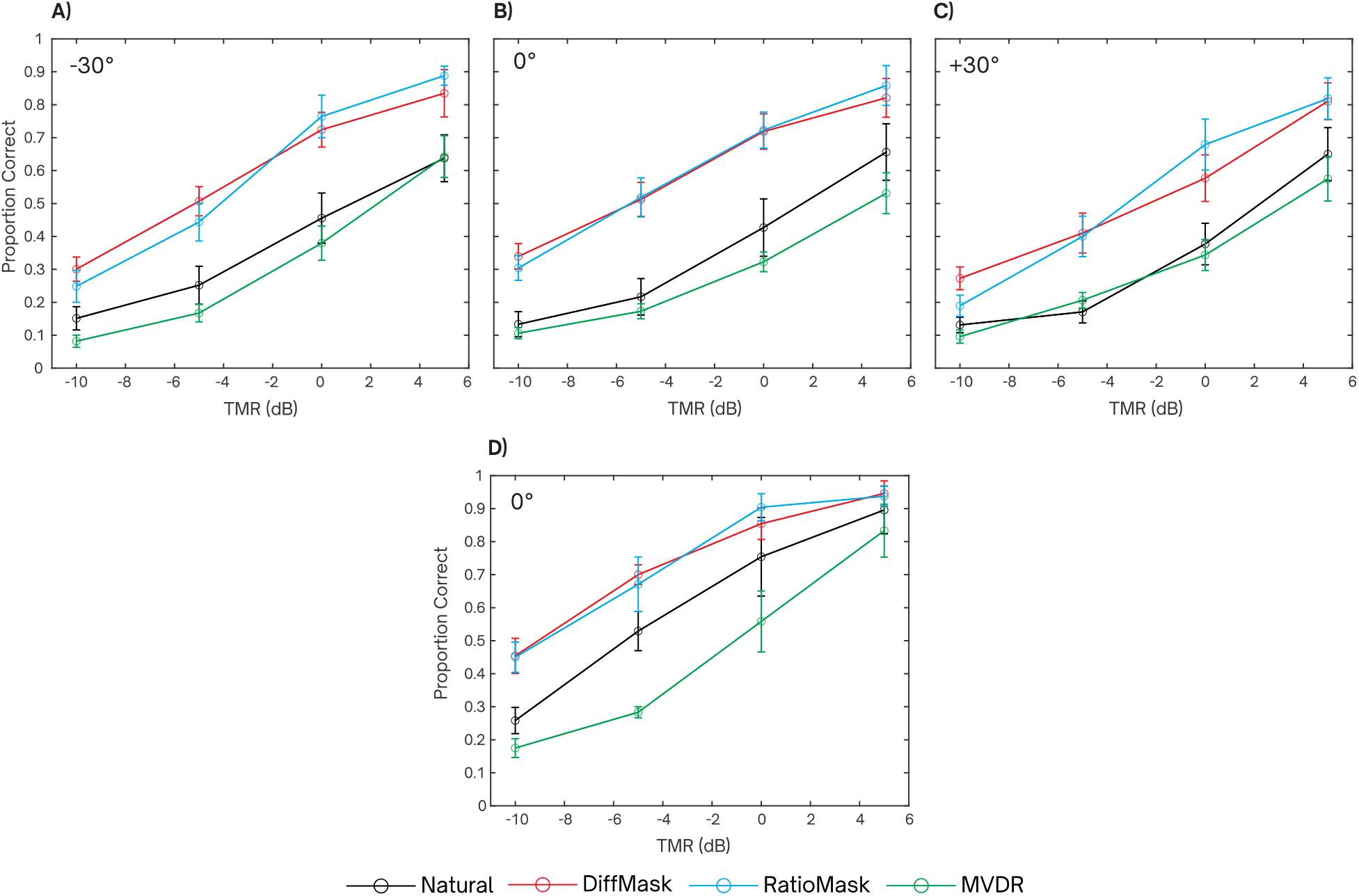
A-C) Average proportion of correctly recognized words as a function of TMR for Experiment 1 (n=8) with target locations of -30°, 0°, +30° respectively. D) Average proportion of correctly recognized words as a function of TMR for Experiment 2 (n = 4) with a target location of 0°. Error bars show across-subject standard deviations.

These patterns are summarized in Figure 2, which shows group-mean speech reception thresholds (SRTs) for each processing condition in each experiment. For each experiment, a one-way repeated measures ANOVA found a significant main effect of processing condition (Exp 1A [F(3,21)=50.77, p<0.001], Exp 1B [F(3,21)=41.22, p<0.001], Exp 1C [F(3,21)=106.74, p<0.001], Exp 2 [F(3,9)=29.43, p=<0.001]). Planned comparisons (paired t-tests, p<0.01) indicated that for Experiments 1A,1B, and 2, both versions of BOSSA resulted in better SRTs than the Natural condition, whereas for Experiment 1C this was only true for RatioMask. In all cases, SRTs for MVDR and Natural were not significantly different.

**Figure 2.**
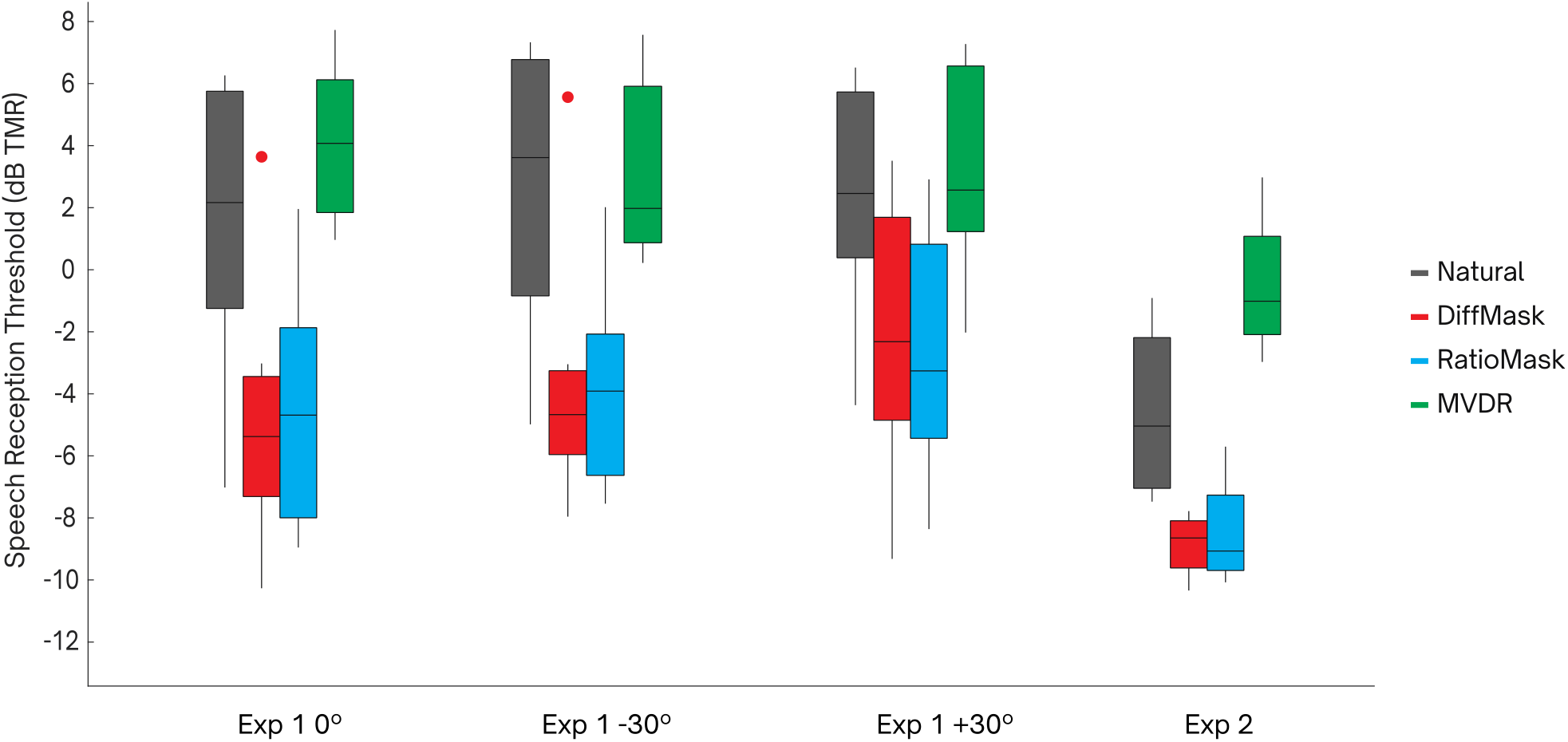
Speech reception thresholds shown as box plots for each processing condition in Experiment 1 (n=8) and Experiment 2 (n=4). Outliers demarcated with a dot exceeded 1.5 IQR.

Figure 3 shows group-mean SRT differences for each of the processing conditions compared to the Natural condition. These differences are expressed in terms of a processing “benefit”, where a positive value in dB corresponds to a decrease in SRT, and a negative value in dB corresponds to an increase in SRT.

**Figure 3.**
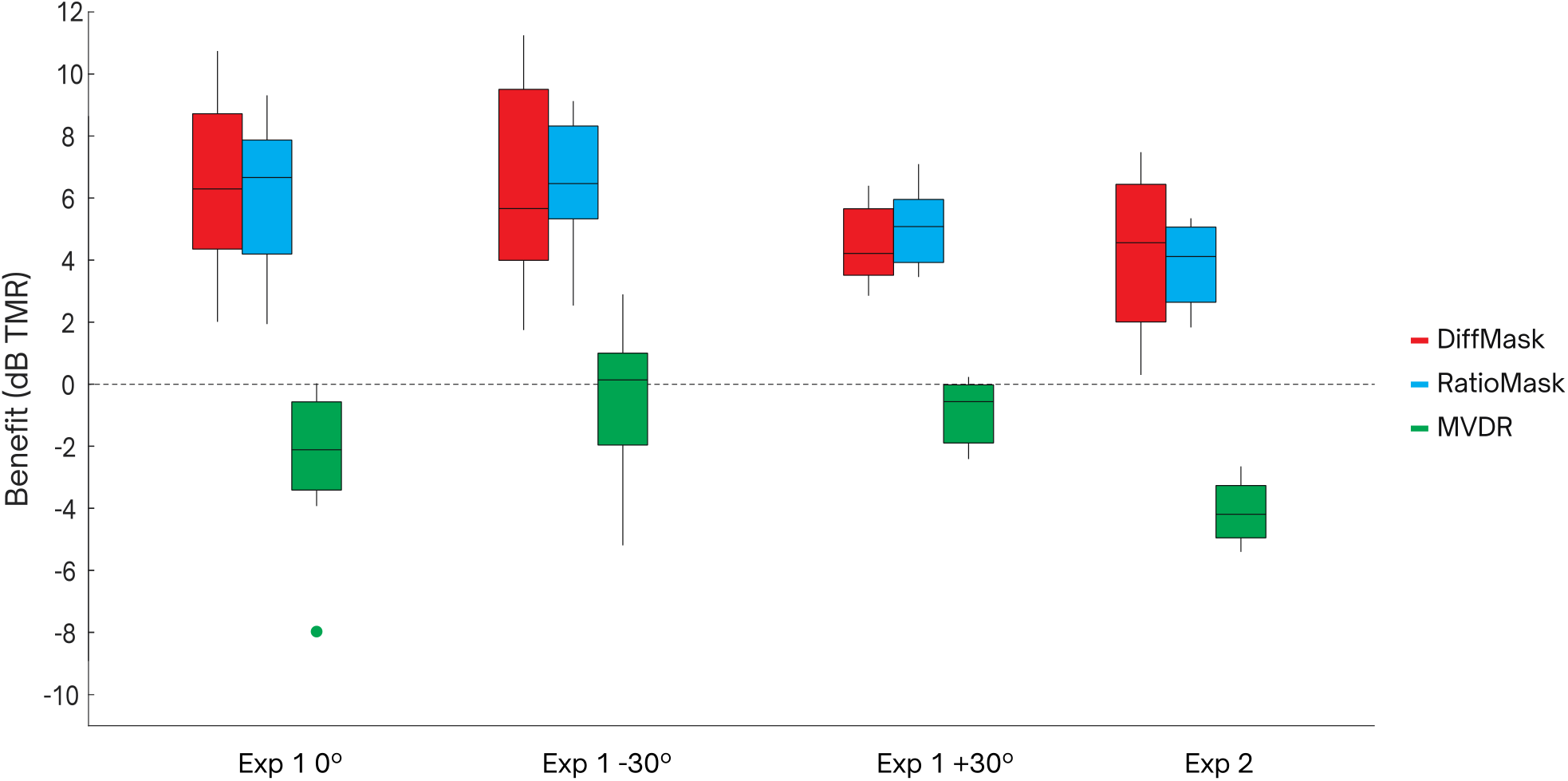
Processing benefits shown as boxplots for each processing condition in Experiment 1 (n=8) and Experiment 2 (n=4). Outliers demarcated with a dot exceeded 1.5 IQR.

## DISCUSSION

### Benefits of BOSSA in Relation to Natural Listening and Beamforming

We have developed a brain-inspired algorithm (BOSSA) that can selectively enhance one talker in a multitalker mixture according to its spatial location. The algorithm has a wide variety of potential uses, including in hearing-assistive devices, where it could provide a benefit in challenging communication situations to millions of people with hearing difficulties. In the current study we evaluated BOSSA in adults with bilateral sensorineural hearing loss and compared its performance to a standard beamforming approach used in hearing aids.

Our evaluation used a challenging speech mixture consisting of five female talkers uttering sentences that were highly confusable with each other. This amounted to a speech intelligibility task with extremely high “informational masking.”^2^ We considered two versions of this task (Experiments 1 and 2) to confirm that the results were not dependent on specific details of the task, and we considered three different target locations, motivated by the idea that a listener may wish to listen to a target that is not directly in front and an assistive algorithm may need to be “steered” to that location. BOSSA improved speech reception thresholds consistently across these variations, relative to the natural condition with no processing. While the benefit was robust, its magnitude varied across participants and configurations, ranging from 0.3 to 11.3 dB.

Multichannel beamformers like MVDR are designed to provide robust benefits in situations where speech is masked by stationary noise (“energetic masking”; air conditioning noise, etc.) and in such cases can improve speech intelligibility, speech quality, listening effort, and numerous objective measures.^15^ At the outset, it was unclear how the MVDR algorithm would perform in our challenging multitalker scenario. We found that this approach did not provide a significant benefit, identifying a potential factor in the failure of current hearing aids to provide robust benefits in cocktail party settings. To confirm our binaural implementation of MVDR was working as expected, we conducted a control experiment using a more traditional speech-in-noise design. For this experiment, we brought back four participants from Experiment 1 (the same four who completed Experiment 2). The target was identical to the 0° target from Experiment 1, and the maskers were two independent speech-shaped noises positioned at -90° and +90° azimuth. The spectrum of these noises was matched to the long-term average spectrum of the set of female talkers in the corpus, and the noises were matched in length to the target on each trial. A repeated-measures ANOVA found a significant effect of processing condition [F(3,9)=36.69, p<0.001]. Planned comparisons showed performance for MVDR was slightly better than for the natural condition (p=0.041), confirming that MVDR was working as expected (Supplemental Figure 1). The performance for the two variations of BOSSA was slightly lower than the natural condition for these stimuli (DiffMask p=0.002, RatioMask p=0.004), confirming that the algorithm in its current form performs best in conditions involving fluctuating maskers. Future versions of BOSSA that are optimized for stationary noise conditions, and/or which incorporate postprocessing like that commonly found in beamforming, may improve the performance of BOSSA under these conditions.

### Comparison To Other Sound Source Segregation Approaches

BOSSA is not the first algorithm to leverage binaural cues to isolate a sound of interest. Roman et al^16^ proposed a related mask-based approach that uses supervised learning, applied to specific scenarios and within frequency bands, to estimate a binary mask from distributions of interaural time and level differences. While this approach is not highly practical for a hearing-aid application due to the complexity of its mask calculation and reliance on training data, it provides robust SNR improvements, increases in automatic speech recognition accuracy, and showed intelligibility improvements in NH listeners under some conditions. The approach was later extended to reverberant situations, with good results according to objective metrics, but no intelligibility data were presented.^17^ To our knowledge, that particular approach has never been evaluated in HI listeners.

Another class of algorithms use much larger numbers of microphones to achieve spatial filtering. To give one example, Kidd and colleagues^18^ developed a hybrid beamformer (“BEAMAR”), which combines the output of a 16-channel beamforming microphone array with natural low-frequency cues to preserve spatial information. Best et al^9^ tested this beamformer for a frontal speech target against four symmetrically separated maskers (a very similar set up to the current study) and produced robust benefits for NH and HI populations. A direct comparison of BOSSA and BEAMAR in a new group of NH listeners^14^ confirmed that the benefits were comparable for the two methods, despite BOSSA using only two microphones compared to 16 microphones for BEAMAR.

In their large comparative study, Volker et al^19^ fitted NH and HI participants with a hearing-aid simulator programmed to run eight state-of-the-art “pre-processing strategies” that included binaural algorithms as well as several single- and multi-channel noise-reduction algorithms. SRT benefits were broadly similar across strategies (on the order of 2-5 dB across three different background noises). The current study suggests that BOSSA would outperform these state-of-the-art strategies for listening scenarios containing competing talkers and substantial amounts of informational masking. Future work is needed, however, to comprehensively test BOSSA in a wide variety of listening scenarios and in a larger and more diverse listener population.

Deep neural network (DNN) approaches to sound source segregation are rapidly evolving. In a recent review of their single-channel DNN-based noise reduction strategy, Healy et al^11^ point out the large strides that have been made since its inception a decade ago, both in terms of efficacy and viability for real-time implementation. Similar results are emerging from other groups^12^ and many of the major hearing-aid manufacturers are now incorporating DNN-based noise reduction into their premium hearing devices.^20,21^ It is worth noting that although these approaches achieve impressive results for speech in noise, they remain challenged by competing talkers, an important real-world scenario for listeners. Another challenge for DNN-based approaches is the requirement of large labeled datasets for training. Generating such a dataset is labor intensive and potentially costly. Moreover, even when such training datasets are available, generalization to scenarios that are not part of the training has not been convincingly demonstrated even for state-of-the-art DNNs. Part of the issue here is that it is difficult to thoroughly sample the vast number of possible configurations of target and maskers in a complex scene with multiple sources in a training dataset. Finally, despite their impressive performance under many conditions, the power consumption required for the highly intensive computational demands of DNNs is also a factor that continues to impact their adoption in miniature wearable devices such as hearing aids. In this context, it is worth noting that the neurally inspired architecture of BOSSA makes it well suited for power-efficient neuromorphic implementations.

### Limitations and Future Work

While BOSSA provided robust improvements in speech intelligibility under the conditions of our experiments, there are current limitations that could hinder its performance in real-world scenarios. For example, in this evaluation, the number and location of spatially tuned neurons (STNs) were fixed according to the known locations of the competing talkers in the mixture. Further work is needed to develop and optimize versions of BOSSA with a dynamic set of STNs capable of selectively isolating sounds from any direction given any arbitrary mixture of competitor locations. This could be accomplished by decoding the neural activity of STNs to first perform source localization and prioritization before proceeding to source isolation.

Computational headroom currently limits the number of STNs that contribute to the reconstruction mask calculation with acceptable latency, but intelligent approaches for selecting the ideal set of STNs for reconstruction may yield robust real-world performance while maintaining the algorithm’s high efficiency. While we have favored biological interpretability over deep-learning approaches in this study, such techniques may be helpful in guiding the optimization of BOSSA for a wider range of acoustic scenes than those tested here. In addition, future implementations of BOSSA will take input directly from the user to dynamically change its focus instead of assuming a fixed look direction. One such method that is actively being explored is to use eye gaze patterns as an input source, creating a means for informed steering or tuning of the spatial filter. Behavioral studies using such an approach suggest that listeners with and without hearing loss are able to use and gain speech intelligibility benefits from this kind of system.^22,23,24^

## METHODS

### Participants

Participants in the experiments were eight adults (ages 20-42 years) with bilateral sensorineural hearing loss. Their hearing losses represented a wide variety of configurations and severities but were all bilaterally symmetric; across-ear average audiograms are shown in Figure 4.

**Figure 4.**
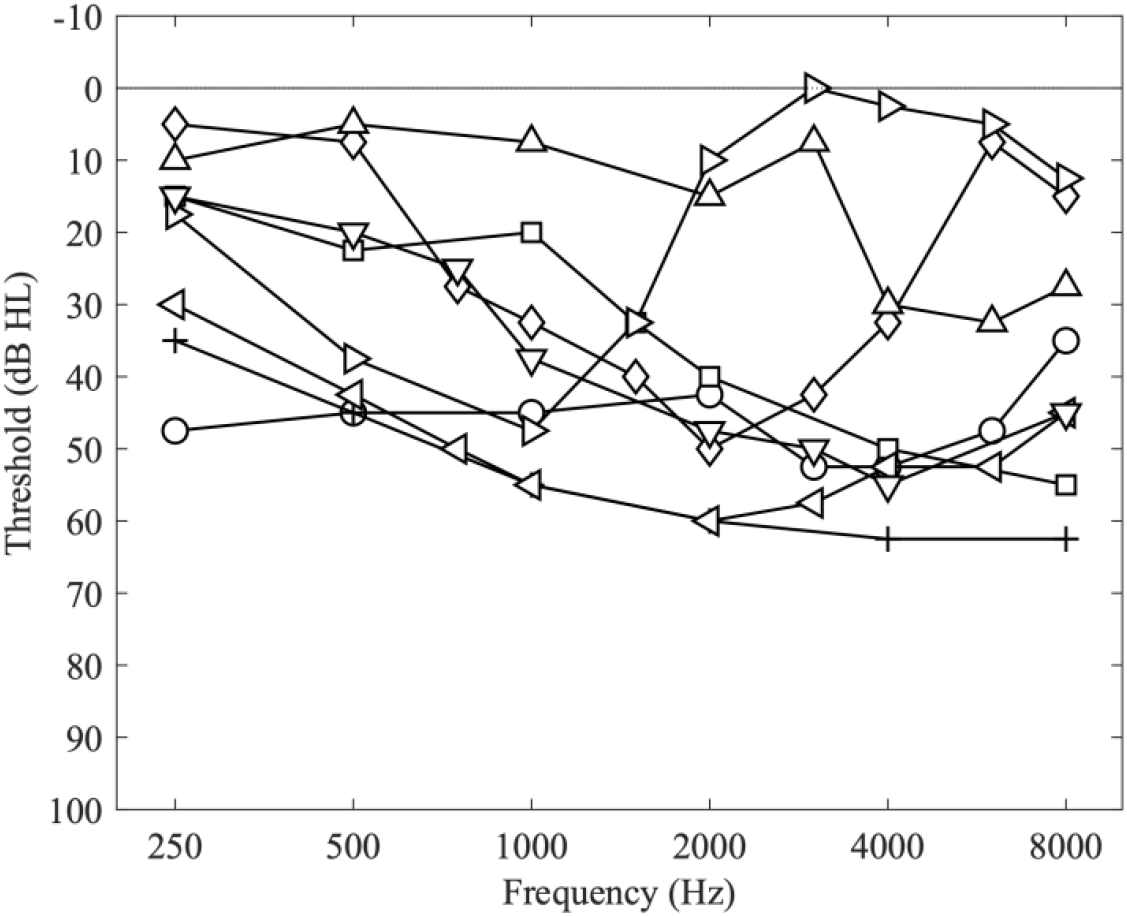
Individual audiograms (averaged over left and right ears) for each participant.

Participants were paid for their participation and gave written informed consent. All procedures were approved by the Boston University Institutional Review Board. All participants completed Experiment 1, and a subset of four participants completed Experiment 2.

### Stimuli

Five-word sentences were constructed from a corpus of monosyllabic words^25^ with the form [name-verb-number-adjective-noun] (e.g., “Sue found three red hats”). The corpus contains eight words in each of the five categories. On each trial, a target sentence was mixed with four masker sentences. The target sentence was designated by the name “Sue”. The five sentences were simulated to originate from five spatial locations (0°, ±30°, and ±60° azimuth) by convolving with anechoic head-related transfer functions measured on an acoustic manikin^26^ at a distance of 1m. The level of the target was varied to achieve target-to-masker ratios (TMRs) of –10, -5, 0, and 5 dB. The nominal presentation level of the mixture (post-processing, see below) was 62 dB SPL. On top of this, each listener was given linear frequency-dependent gain to compensate for their audiogram according to the NAL-RP prescriptive formula.^6^

In Experiment 1 (Figure 5A), each word in a sentence was spoken by a different female talker, randomly chosen from a set of eight female talkers, without repetition. The target and masker sentences were time-aligned, so that the five words in each category shared the same onset, and zero-padding was applied to all but the longest word in each category to align their offsets. The design of these stimuli was intended to reduce the availability of voice and timing-related cues, and as such increase the listener’s reliance on spatial information to solve the task. In three separate sub-experiments, the target sentence was presented from 0°, -30°, and +30° azimuth, and the four maskers were presented from the other four non-target locations in the set.

**Figure 5.**
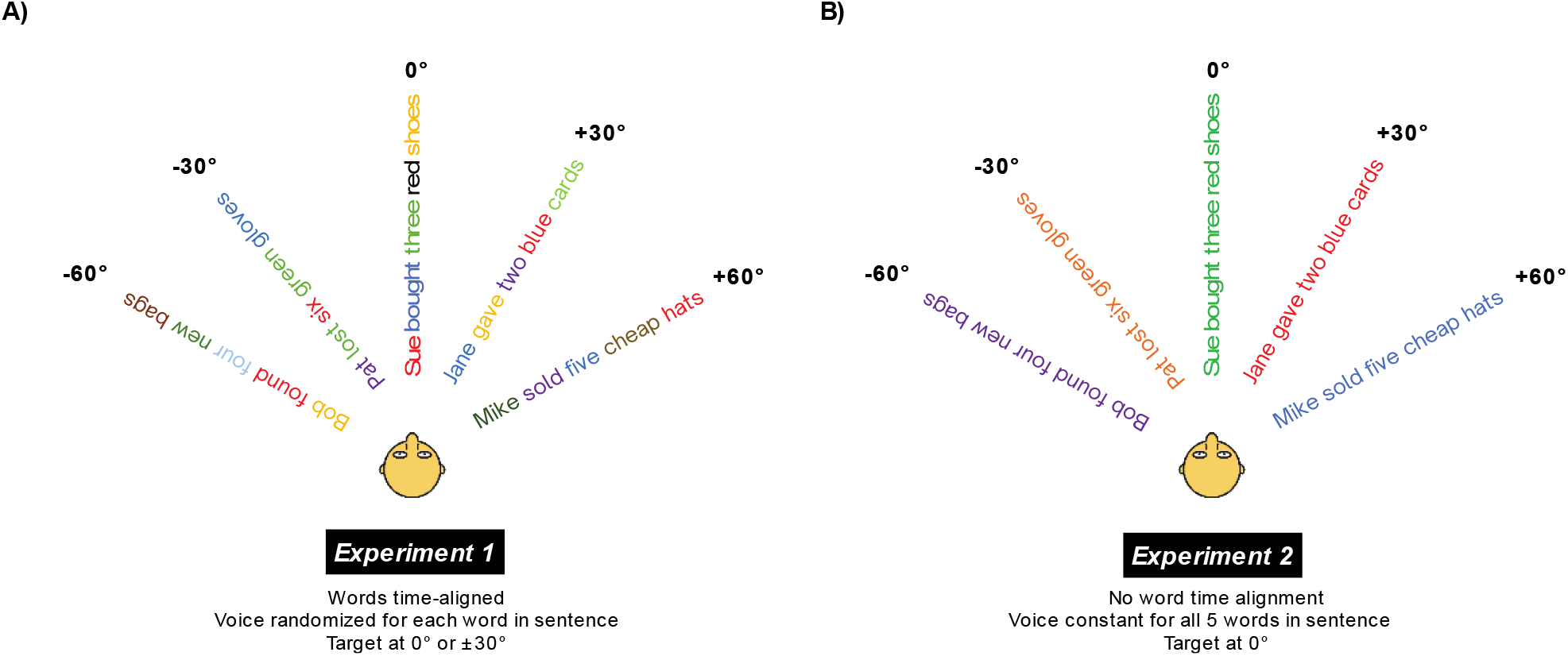
A) Talker structure and spatial configuration of the five competing sentences in Experiment 1. The target source locations tested were 0°, -30°, and +30° azimuth. B) Talker structure and spatial configuration of the five competing sentences in Experiment 2. The target source location tested was 0°. Different colors depict different female talkers.

In Experiment 2 (Figure 5B), two modifications were made to ensure that the results were not dependent on specific choices made in Experiment 1. In this experiment, the words in each sentence were spoken by the same talker, such that there was voice continuity as well as spatial continuity within each competing sentence. In addition, the onset of words were not time-aligned across sentences, but individual word recordings were simply concatenated (with no additional gaps) to create each sentence. For Experiment 2, only the center target location was examined.

### Procedures

Each experiment was comprised of 12 blocks of trials (three blocks for each of the four processing conditions, see below). Each block contained five trials at each of the four TMRs (20 total trials per block). The order of presentation of TMRs within a block, and the order of blocks for each participant, were chosen at random. Each experiment took approximately one hour to complete.

Stimuli were controlled in MATLAB 2019a (MathWorks Inc., Natick, MA) and presented at 44.1 kHz sampling rate through an RME HDSP 9632 24-bit soundcard (RME Audio, Bayern, Germany) to Sennheiser HD280 Pro headphones (Sennheiser Electronic GmbH & Co., Wedemark, Germany). The sound system was calibrated at the headphones with a sound meter (type 2250; Brüel & Kjær, Nærum, Denmark). Participants were seated in a double-walled sound-treated booth. A computer monitor inside the booth displayed a graphical user interface containing a grid of 40 words (five columns of eight words, each column corresponding to one word category). On each trial, participants were presented with a mixture and were instructed to listen for the target sentence. Responses were provided by choosing one word from each column on the grid with a computer mouse. The stimulus set up and the target location (left, center, right) was described to the participant prior to each experiment.

Performance was evaluated by calculating the percentage of correctly answered keywords across all trials at each TMR. Psychometric functions were generated by plotting the percent correct as a function of TMR and fitting a logistic function to those data. SRTs, defined as the TMRs corresponding to 50% correct, were extracted from each function using the psignifit toolbox.^27^ Statistical analysis was performed using IBM SPSS Statistics (Version 29).

### Processing Conditions

The four processing conditions included one control condition that simulated the natural listening condition (“Natural”), two variations of BOSSA (“DiffMask” and “RatioMask”), and a standard binaural beamformer (“MVDR”).

#### BOSSA

Using an approach inspired by ideal time-frequency mask estimation,^28^ BOSSA separates an incoming binaural audio signal into time-frequency bins then selectively applies gain to bins such that sound energy arising from a prescribed target location is preserved while sound energy from non-target locations is suppressed. As described in Chou et al ^14^, the gain for each time-frequency bin was calculated by combining the spiking activity of five sets of STNs with target angles *θ* ∈ {0°, ±30°, ±60°}.

Two versions of BOSSA were evaluated. These algorithms differed only in the content of the reconstruction module responsible for converting ensembles of neural spikes into the final binaural audio output as depicted in Figure 6.

**Figure 6.**
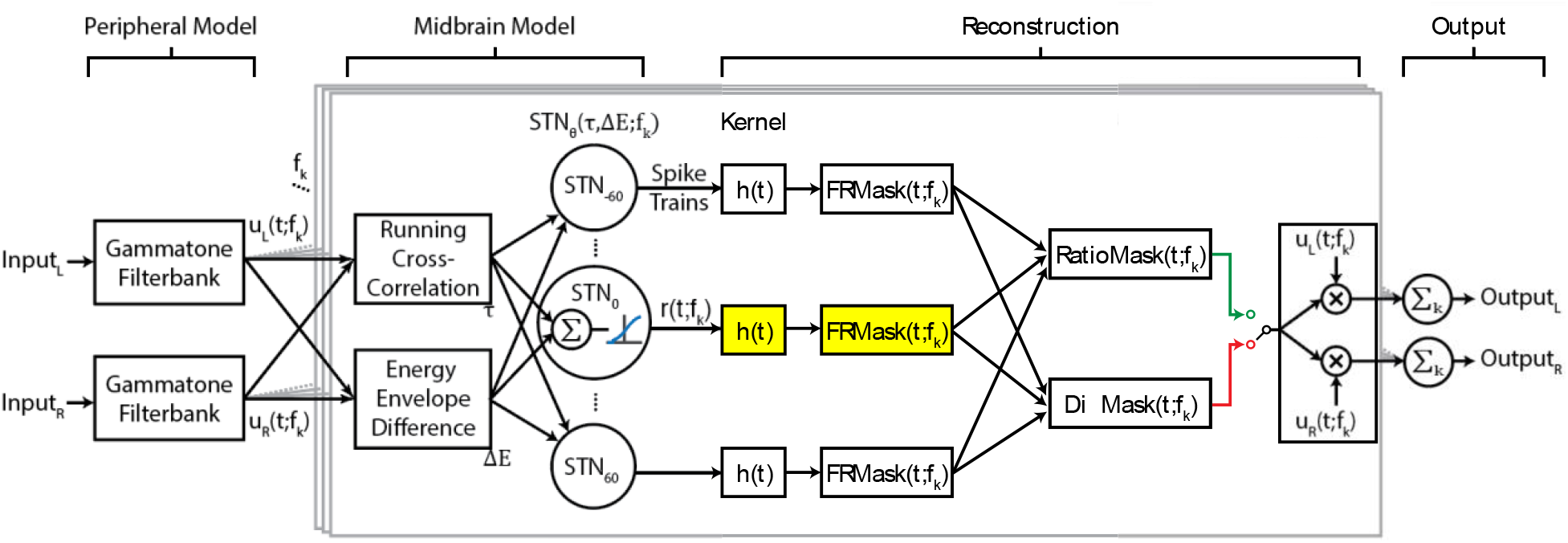
Flow diagram of BOSSA algorithm. Central boxes, outlined in grey, show processing for a single frequency band. The functions *u*_*L*_(*t*; *f*_*k*_) and *u*_*L*_(*t*; *f*_*k*_) are the narrowband signals of the left and right input channels for each frequency channel, and *f*_*k*_ denotes the *k*^*th*^ frequency channel. The midbrain model is based on spatially tuned neurons (STNs), where each STN has a “best” ITD and ILD, denoted *τ* and *ΔE*, respectively. The best ITD and ILD values of a neuron depend on the direction *θ* and frequency *f*_*k*_ to which the STN is tuned. *h*(*t*) represents the reconstruction kernel that converts spike trains to waveforms. Either RatioMask (green line) or DiffMask (red line) was used for reconstruction, as indicated by the switch. The target STN for reconstruction (yellow highlight) is manually selected as a parameter. The implementations of DiffMask and RatioMask in our analysis involves five sets of STNs, where *θ* ∈ {0, ±30, ±60}; however, other implementations of the model may involve different sets of *θ*.S

The first version of BOSSA used a previously studied reconstruction method, DiffMask, which was inspired by lateral inhibition observed in biological networks. In this technique, the scaled sum of non-target STN firing rates 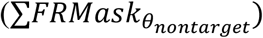 was subtracted from the target STN firing rate activity 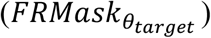. A lower limit of 0 was imposed on the DiffMask output to prevent unrealistic negative firing rates. The subtractive term acts as a mechanism for sharpening the spatial tuning of output neurons as demonstrated in a previous publication.^14^ The scale factor *a* was adjusted to normalize the mask to a maximum value of 1.

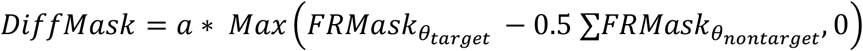

The second version of BOSSA tested in this study utilized a new reconstruction method called RatioMask, intended to reduce some unnatural artifacts induced by BOSSA with DiffMask.

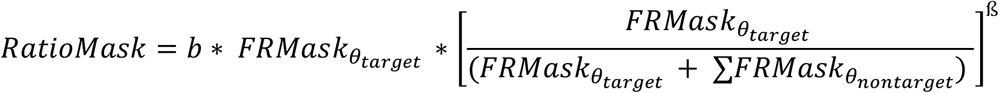

In contrast to DiffMask, which employs a subtractive operation, RatioMask implements a multiplicative operation to sharpen the spatial tuning of output neurons in the presence of competing noise. The multiplicative term, inspired by the ideal ratio mask (IRM) operation,^28^ is an estimate of the SNR raised to a power β. Here, the firing rate of the neuron at the target location serves as an estimate of the “signal” and the firing rates at non-target locations serves as an estimate of the “noise”. Thus, the multiplicative term boosts time-frequency tiles with a higher SNR and suppresses time frequency tiles with lower SNR. We found that a value of β=1.65 gave the best results as quantified by an objective intelligibility metric (see below). The normalization scale factor b was adjusted to yield a maximum value of 1 for the RatioMask.

After a given mask calculation, the mask was then applied (i.e., point-multiplied) to the left and right channels of the original sound mixture. Lastly, the sum of each frequency channel of the mask filtered signal was taken to obtain an audible, segregated waveform. In contrast to DiffMask, RatioMask applies a gain factor without any “hard” thresholding operation, which led to a smoother, more natural sounding output based on our listening experience.

#### BOSSA Parameters

Most model parameters were fixed to biologically plausible values for the implementation of BOSSA evaluated here, rather than chosen based on an extensive optimization process. Variation in some reconstruction parameters, however, was explored using the Short Time Objective Intelligibility (STOI) measure.^29^ STOI ranges between 0 and 1 and can be used to predict speech intelligibility when combined with an appropriate mapping function. The time-constant of the alpha function kernel (τ_h_), and the scaling factor for DiffMask (*a*) (see Chou et al^14^ for details) were chosen by iteratively trying a range of values, quantifying algorithm performance using STOI, then choosing parameter values that produced the highest average STOI. For RatioMask, the beta (ß) parameter value of 1.65 was selected using the genetic algorithm (GA) function in the MATLAB Global Optimization Toolbox with “fitness” defined as the average STOI value.

While we do not claim that either approach outputs the optimal parameter set for reconstruction, our experience and the behavioral results indicate the chosen parameter values produced adequate reconstructions that supported robust speech intelligibility.

#### Binaural MVDR

To compare to a widely used spatial processing algorithm, stimuli were also processed with a binaural MVDR beamformer.^8,30^ To enable a direct comparison to the two-channel BOSSA approach, the binaural MVDR implementation used two virtual microphones (one per ear). Relative transfer function vectors aimed towards the target source angle were calculated for each ear using the same KEMAR HRTFs that were used in BOSSA. Log-level voice activity detection was performed for each ear along with the MVDR application followed by a multichannel Wiener filter with the decision directed approach.^31^

## Acknowledgements

This work was supported by grants from the National Institutes of Health (Award No. DC013286), the National Science Foundation (Award No. 9500314916) and the Demant Foundation. The authors would like to thank Sergi Rotger-Griful, Martin Skogland, Paol Hoang and Gerald Kidd for their input and support as well as Kenny Chou for his prior work on the BOSSA algorithm.

## Author contributions statement

AB and VB designed the psychophysical experiment, conducted the experiment, analyzed the data, and wrote the first version of the manuscript, with editing by KS. AB designed the algorithm under the supervision of KS. All authors contributed to the article and approved the submitted version.

## Data Availability Statement

The raw data supporting the conclusions of this article will be made available by the authors, without undue reservation.

