## Supplemental Figure 1 for "A brain-inspired algorithm improves “cocktail party” listening for individuals with hearing loss"

### Supplementary Information

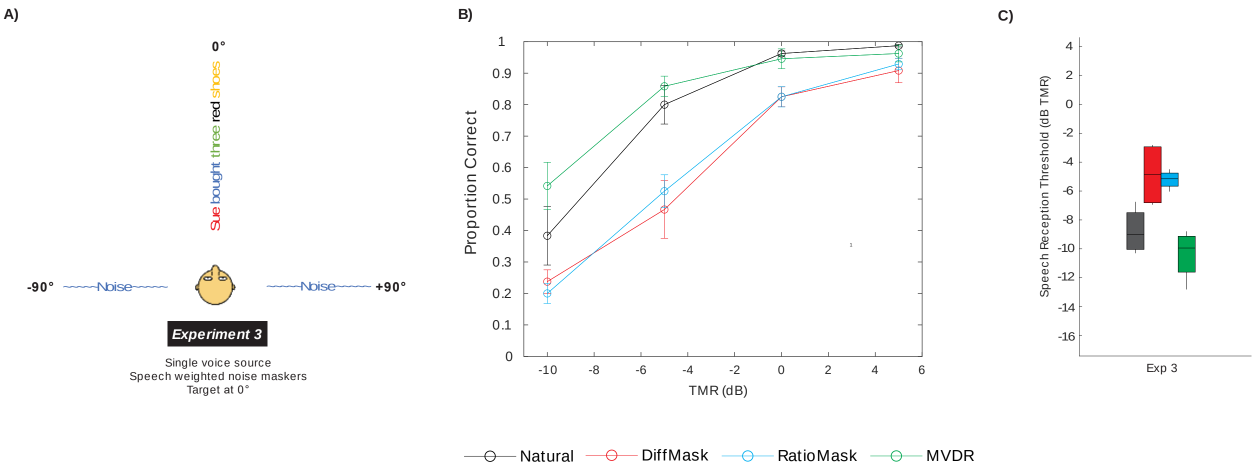


**Figure S1.** A) Talker structure and spatial configuration of the three competing sources in Experiment 3. The target source location tested was 0° and speech weighted noise maskers were positioned at +90˚ and -90˚. B) Average proportion of correctly recognized words as a function of TMR for Experiment 3 (n=4) with a target location of 0˚. Error bars show across-subject standard deviations. C) Speech reception thresholds shown as box plots for each processing condition in Experiment 3.
